# Prediction of RNA-protein interactions using a nucleotide language model

**DOI:** 10.1101/2021.04.27.441365

**Authors:** Keisuke Yamada, Michiaki Hamada

## Abstract

**Motivation:** The accumulation of sequencing data has enabled researchers to predict the interactions between RNA sequences and RNA-binding proteins (RBPs) using novel machine learning techniques. However, existing models are often difficult to interpret and require additional information to sequences. Bidirectional encoder representations from Transformer (BERT) is a language-based deep learning model that is highly interpretable. Therefore, a model based on BERT architecture can potentially overcome such limitations.

**Results:** Here, we propose BERT-RBP as a model to predict RNA-RBP interactions by adapting the BERT architecture pre-trained on a human reference genome. Our model outperformed state-of-the-art prediction models using the eCLIP-seq data of 154 RBPs. The detailed analysis further revealed that BERT-RBP could recognize both the transcript region type and RNA secondary structure only from sequence information. Overall, the results provide insights into the fine-tuning mechanism of BERT in biological contexts and provide evidence of the applicability of the model to other RNA-related problems.

**Availability:** Python source codes are freely available at https://github.com/kkyamada/bert-rbp.

## 1 Introduction

Interactions between RNA sequences and RNA-binding proteins (RBPs) have a wide variety of roles in regulating cellular functions, including mRNA modification, splicing, translation, and localization (Hentze *et al.*, 2018). For instance, the T-cell-restricted intracellular antigen (TIA) family of proteins functions as alternative splicing regulators (Wang *et al.*, 2010), and heterogeneous nuclear ribonucleoprotein K (hnRNPK) is a versatile regulator of RNA metabolism (Geuens *et al.*, 2016). Numerous attempts have been made to identify RNA-RBP interactions to accurately capture their biological roles.

Among the various *in vivo* experimental methods, high-throughput sequencing of RNA isolated by crosslinking immunoprecipitation (CLIP-seq) is widely used to reveal a comprehensive picture of RNA-RBP interactions (Licatalosi *et al.*, 2008; Lin and Miles, 2019). Altered CLIP-seq protocols have also been developed (Hafner *et al.*, 2010; König *et al.*, 2010; Van Nostrand *et al.*, 2016). Recently, a large amount of enhanced CLIP-seq (eCLIP) data, targeting more than 150 different RBPs, was generated during phase III of the Encyclopedia of DNA Elements (ENCODE) Project (Van Nostrand *et al.*, 2020).

Because there is a vast volume of available CLIP-seq data, recent bioinformatics studies have focused on developing machine learning models to predict RNA-RBP interactions and deciphering hidden patterns translated by these models (Pan *et al.*, 2019; Yan and Zhu, 2020). Early models use statistical evaluations or support vector machines (SVMs) to classify RNA sequences into RBP-bound or RBP-unbound groups (Hiller *et al.*, 2006; Kazan *et al.*, 2010; Maticzka *et al.*, 2014). One of the SVM-based models, GraphProt, encodes RNA sequences and their estimated secondary structures into graph representations (Maticzka *et al.*, 2014). Non-negative matrix factorization (NMF) and random forest were also adapted to other models (Stražar *et al.*, 2016; Yu *et al.*, 2019). Since Alipanahi *et al.* (2015) demonstrated the applicability of convolutional neural networks (CNNs) for predicting RNA-protein and DNA-protein interactions, several deep learning models have been developed. While some models incorporate a single CNN with some modifications (Pan and Shen, 2018; Zhang *et al.*, 2019; Tahir *et al.*, 2021), others use a different neural network model (Uhl *et al.*, 2020) or a combination of several neural network architectures (Ben-Bassat *et al.*, 2018; Pan *et al.*, 2018; Yan *et al.*, 2020; Deng *et al.*, 2020; Grønning *et al.*, 2020). For instance, HOCNNLB uses high-order encodings of RNA sequences as inputs for CNN (Zhang *et al.*, 2019), and iDeepS uses stacked CNN and bidirectional long short-term memory (biLSTM) and takes both RNA sequences and their estimated secondary structures as inputs (Pan *et al.*, 2018). However, existing models often lack interpretability because of the complex nature of the neural network and require additional information to RNA sequences. Therefore, the development of advanced models that overcome these limitations is awaited.

The improvement of deep learning architectures largely buttresses progress in building better bioinformatics tools. In the field of natural language processing, self-attention-based deep learning architectures, such as Transformer and BERT, have achieved state-of-the-art performance in various tasks (Vaswani *et al.*, 2017; Devlin *et al.*, 2019). Additionally, BERT, which essentially consists of stacked Transformer encoder layers, shows enhanced performance in down-stream task-specific predictions after pre-training on a massive dataset (Devlin *et al.*, 2019). In the field of bioinformatics, several BERT architectures pre-trained on a massive corpus of protein sequences have been recently proposed, demonstrating their capability to decode the context of biological sequences (Rao *et al.*, 2019; Rives *et al.*, 2021; Elnaggar *et al.*, 2021; Iuchi *et al.*, 2021). In comparison to the protein language models, Ji *et al.* (2021) a pre-trained BERT model, named DNABERT, on a whole human reference genome demonstrated its broad applicability for predicting promoter regions, splicing sites, and transcription factor binding sites upon fine-tuning. Thus, pre-trained BERT models are potentially advantageous for a wide variety of bioinformatics tasks, including the prediction of RNA-RBP interactions.

In addition to its performance, BERT is highly interpretable and suitable for translating extended contextual information compared to conventional deep learning architectures, such as CNNs and long short-term memory (Rogers *et al.*, 2020). Researchers in an emerging field, called BERTology, intend to elucidate how BERT learns contextual information by analyzing attention, which essentially represents the flow of information within a model (Vig and Belinkov, 2019). For instance, analysis of protein BERT models revealed that protein contact maps could be reconstructed from the attention of the model (Vig *et al.*, 2021; Rao *et al.*, 2021). This implies that, by analyzing the fine-tuned BERT model, we can reasonably explain the types of features that are crucial for predicting RNA-RBP interactions.

In this study, we applied the BERT model pre-trained on a human reference genome to predict the RBP-binding property of RNA sequences. Our model, named BERT-RBP, outperformed existing state-of-the-art models as well as the baseline BERT model whose weight parameters were randomly initialized, showing the significance of pre-training on a large corpus. Attention analysis on the fine-tuned model further revealed that BERT-RBP could translate biological contexts, such as transcript region type, transcript region boundary, and RNA secondary structure, only from RNA sequences. Thus, this study highlights the powerful capability of BERT in predicting RNA-RBP interactions and provides evidence of the architecture’s potential applicability to other bioinformatics problems.

## 2 Materials and methods

### 2.1 Terminology

#### k-mer

For a given sequence, k-mers of the sequence consisted of every possible subsequence with length k, i.e., given a sequence *ACGTAC*, the 3-mers of this sequence included *ACG*, *CGT*, *GTA*, and *TAC*, and the 4-mers included *ACGT*, *CGTA*, and *GTAC*.

#### Token

Tokens are referred to as words or their positions within a sequence. Tokens included not only k-mers but also special tokens, such as CLS (classification) and SEP (separation). In our model, CLS was appended at the beginning of each input sequence, and its feature vector from the final layer was used for classification. The SEP was attached only to the end of each sequence.

#### Attention

Attention represents the flow of information within the BERT model. The attention weight indicates how much information the hidden state of a token in the upper (closer to the output) layer referred to the hidden state of a token in the lower (closer to the input) layer.

### 2.2 Data Preparation

An eCLIP-seq dataset previously generated from the ENCODE3 database by Pan *et al.* (2020) was used. The original dataset consisted of 154 RBP sets with up to 60,000 positive RNA sequences that bind to the corresponding RBP and the same number of negative sequences. Each positive sequence had a length of 101 nucleotides with the eCLIP-seq read peak at its center, while each negative sequence was sampled from the non-peak region of the same reference transcript as its positive counterpart. First, all sequences that included unannotated regions or repeatedly appeared in the same set were removed to create the dataset, and 15,000 positive and negative sequences were randomly sampled. If the original RBP set included less than 15,000 samples, all sequences were retrieved. The resulting samples were split into training (64%), evaluation (16%), and test sets (20%). For the selected RBPs, non-training datasets were created that included all positive and negative samples, except those in the training sets. The training and evaluation sets were used during the fine-tuning step and cross-validation, the individual test sets were used to measure performance after fine-tuning, and the non-training sets were used for analysis. Additionally, to evaluate models using datasets that include sequences distinct from each other, we applied MMseqs2 (Steinegger and Söding, 2017), which clusters sequence sets into groups of similar sequences, to each original RBP set with the maximum E-value threshold 0.001. After removing redundant sequences from the original dataset, the same number of sequences were sampled as training and evaluation sets to create non-redundant datasets for cross-validation.

### 2.3 Models and Training

#### 2.3.1 Pre-trained BERT model

DNABERT, a BERT-based architecture pre-trained on a human reference genome, was adapted to model RNA sequences and their RBP-binding properties. The parameters of DNABERT were transferred to our BERT models and used to initialize them. Briefly, DNABERT was pre-trained by Ji *et al.* (2021) using k-mer (k = 3-6) representations of nucleotide sequences obtained from a human reference genome, GRCh38.p13. Once the CLS and SEP tokens were appended to the input k-mers, each token was embedded into real vectors with 768 dimensions. The model was pre-trained with the masked language modeling objective, self-supervised learning, to predict randomly masked tokens using information from other tokens. The model had 12 Transformer encoder layers, each of which consisted of 12 self-attention heads and utilized the multi-head self-attention mechanism (Figure 1).

**Figure 1:**
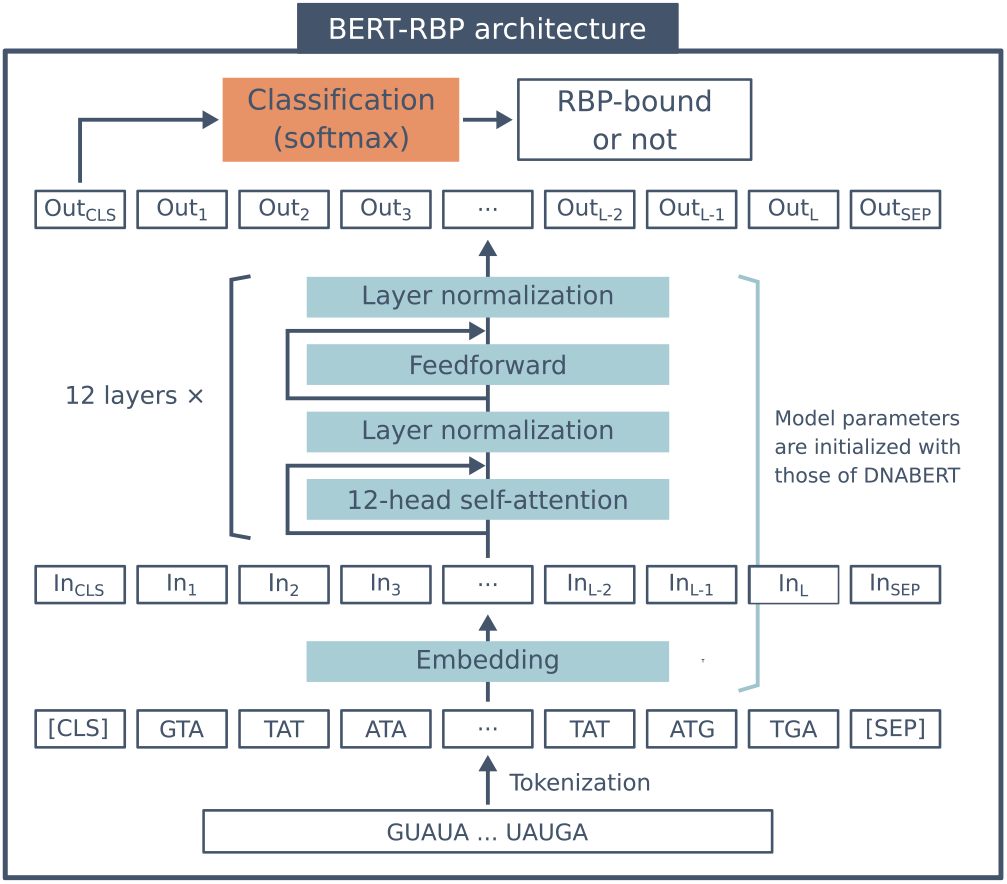
The architecture of BERT-RBP. The input RNA sequence was first tokenized into 3-mers and modified with CLS (classification) and SEP (separation) tokens. Then, each token was embedded into a 768-dimensional feature vector. These feature vectors were consequently processed through 12 Transformer encoder layers, where each layer included 12 self-attention heads. The CLS token of the output vector from the last layer was further utilized for classification to predict whether the input RNA sequence bound to the RBP. Upon fine-tuning, the model parameters of the embedding layer and the stacked Transformer encoder layers were initialized with those of DNABERT. These parameters were randomly initialized for the BERT-baseline model.

#### 2.3.2 Fine-tuning

Upon fine-tuning, the parameters of the model were initialized with those of DNABERT (Figure 1). Subsequently, BERT-RBP was fine-tuned on the training datasets. The hyperparameters used for training are listed in Supplementary Table S1, and these hyperparameters were kept consistent for all the different k-mer models (k = 3-6). The models were trained on four NVIDIA Tesla V100 GPUs (128GB memory). The training of one RBP model using 19,200 samples took less than 10 min. After fine-tuning, the model performance was measured using independent test sets using the area under the receiver operating characteristic curve (AUROC).

#### 2.3.3 Baseline Models

The following four existing models were implemented as baselines: GraphProt, iDeepS, HOCNNLB, and DeepCLIP. GraphProt is an SVM-based model that converts RNA sequences and their estimated secondary structures into graph representations and predicts RBP-binding sites (Maticzka *et al.*, 2014). iDeepS uses a combination of CNN and biLSTM to predict RBP-binding sites from RNA sequences and their estimated secondary structures (Pan *et al.*, 2018). HOCNNLB is another method for training CNNs to predict RBP binding sites while taking k-mer representations of RNA sequences (Zhang *et al.*, 2019). DeepCLIP consists of one convolutional layer that extracts sequence features and following biLSTM to detect RBP binding sites (Grønning *et al.*, 2020). In addition to the above models, the baseline BERT model (BERT-baseline), whose parameters were randomly initialized instead of transferring parameters from DNABERT, was also trained. Hyperparameters used for training BERT-baseline are listed in Supplementary Table S1. All baseline models were trained and tested using the same training and independent test sets as BERT-RBP.

### 2.4 Attention Analysis

We examined whether attention reflected a ny b iological features of the input RNA sequences after fine-tuning. The method proposed by Vig *et al.* (2021) was adapted to ask whether attention agrees with hidden properties of inputs both at the sequence level (transcript region type) and at the token level (RNA secondary structure).

#### 2.4.1 Sequence-level and Token-level Properties

When conditioned by an input sequence, each head emits a set of attention weights *α*, where *α*_*i,j*_ (> 0) indicates the attention weight from the *i*th token in the upper layer to the *j*th token in the lower layer. Σ_*j*_ *α*_*i,j*_ = 1 is satisfied, as the attention weights are normalized over each token in the upper layer. We calculated the attention weights for the CLS in each head as follows:

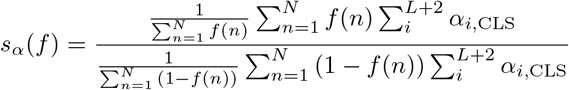

where *N* indicates the number of input sequences; *f* (*n*) is an indicator that returns 1 if the property exists in the *n*th sequence and 0 otherwise; *L* indicates the number of sequence tokens in the *n*th sequence, and *L* + 2 is the number of all tokens, including CLS and SEP. Intuitively, *s*_*α*_(*f*) represents the relative attention to the CLS associated with the property *f*.

For token-level analysis, attention weights to the token of interest were computed at each head using the following equation:

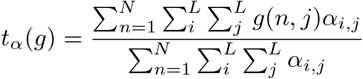

where *g*(*n, j*) is an indicator that returns 1 if the property exists in the *j*th token of the *n*th sequence in the lower layer and 0 otherwise. Note that attention weights to CLS and SEP were not considered during the token-level analysis, and the token length *L* was used. Here, *t*_*α*_(*g*) represents the ratio of attention to property *g*.

#### 2.4.2 Analysis of Transcript Region Type

We first examined whether attention weights reflect transcript region types, including the 5’UTR, 3’UTR, intron, and CDS. Region-type annotations were downloaded from the Ensembl database (Ensembl Genes 103, GRCh38.p13) (Yates *et al.*, 2020). For each gene, we selected the most prominent isoform based on the APPRIS annotation (Rodriguez *et al.*, 2013), transcript support level, and length of the transcript (the longer, the better). The original eCLIP dataset curated by (Pan *et al.*, 2020) used GRCh37/hg19 as a reference genome, so we converted sequence positions into those of GRCh38/hg38 using the UCSC liftOver tool (Kent *et al.*, 2002) and retained those sequences that could be remapped with 100% sequence identity. For simplicity, sequences containing one or more nucleotides labeled with the region type were regarded as having that property. Using the non-training dataset, we accumulated attention weights to the CLS token at each head, averaged over the region type, and calculated the attention level relative to the background (Equation 2.4.1). Finally, the coefficient of variation of the relative attention level among 144 heads was computed to measure the degree of specialization of each BERT model.

#### 2.4.3 Analysis of RNA Secondary Structure

The secondary structure of RNA was another property that was analyzed. For each input RNA sequence, the structure was estimated based on the maximum expected accuracy (MEA) using LinearPartition (Zhang *et al.*, 2020). Once the MEA structures were estimated, each nucleotide was labeled with one of six structural properties; F (dangling start), T (dangling end), I (internal loop), H (hairpin loop), M (multi-branched loop), and S (stem). 3-mer tokens containing one or more nucleotides labeled with structural properties were defined as having the structure. The ratio of attention weights to the structural property (Equation 2.4.1) was computed for each head and compared to the overall probability of the structure within the dataset. The similarity of attention ratio patterns between DNABERT and BERT-RBP was measured using Kendall’s rank correlation coefficient to assess the degree of variation from DNABERT to BERT-RBP. For each RBP, models were fine-tuned using six different seed values to consider the probabilistic behavior during fine-tuning, and the mean of the six correlation coefficients was calculated. The head, which showed the most significant attention ratio, was selected for each RBP and each structure type, and the raw attention weights 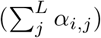 were extracted from the head for each token. Because of the nature of attention, the number of samples tends to be sparse in the range where attention weights are relatively high; therefore, samples whose attention weights were within the 99.5 percentile were the focus. Consequently, we analyzed the relationship between the raw attention weight to the structure and the 3-mer token probability to have the structural property.

#### 2.4.4 Motif Extraction

By optimizing DNABERT’s motif analysis pipeline for the RBP scenario, we extracted RBP binding motifs from BERT-RBP. Using positive samples in the test dataset of each RBP, the attention weights from CLS to each token (*α*_*CLS,j*_) were accumulated over 12 heads of the final layer of BERT-RBP. Extracted attention weights were used to detect consequent high attention regions within each sequence; those high attention regions were restricted to six- to ten-nucleotide long and treated as motif candidates. These candidates were then filtered using the Hypergeometric test with a 0.005 p-value cut-off to declare statistically significant enrichment in the positive samples. Finally, similar motif candidates were merged using pair-wise alignment, and resulting motifs with the highest number of instances were selected. Selected motifs were compared with those detected using an *ab initio* motif discovery tool, STREME (Bailey, 2021), on the same test dataset, and motifs recorded in the mCrossBase (Feng *et al.*, 2019), which is a database of RBP binding motifs and crosslink sites defined by using ENCODE eCLIP data.

## 3 Results

### 3.1 Performance of BERT-RBP

We evaluated the prediction performance of our model along with four existing models (GraphProt, iDeepS, HOCNNLB, and DeepCLIP) and BERT-baseline. All models were trained on the same training set, and their performance was measured using an independent test set over 154 RBPs. BERT-RBP resulted in an average AUROC of 0.786, which was higher than that of any other model (Figure 2). The average AU-ROCs of the other models were 0.691, 0.769, 0.771, 0.680, and 0.767 for GraphProt, iDeepS, HOCNNLB, DeepCLIP, and BERT-baseline, respectively. All scores are reported in the Supplementary Table S2 along with other metrics. While the baseline BERT model alone showed comparable performance to existing methods, our model improved the scores, indicating the significance of pre-training on a large DNA corpus to predict RNA-RBP interactions. In addition, the score of BERT-RBP was higher than the previously reported average AUROC (0.781) of the updated iDeepS (Pan *et al.*, 2020), even though we used approximately five times smaller subsets of their training data to train BERT-RBP. A one-to-one comparison against each baseline revealed that our model demonstrated the best scores for 146 out of 154 RBPs (Supplementary Figure S1). The model performance was also evaluated with five-fold cross-validation using the original training and evaluation sets and using the low-similarity datasets. BERT-RBP showed the best scores for 149 and 152 RBPs for each cross-validation, respectively (Supplementary Figure S2). These results demonstrated that our model exceeded the state-of-the-art methods in predicting RNA-RBP interactions while taking only sequence information.

**Figure 2:**
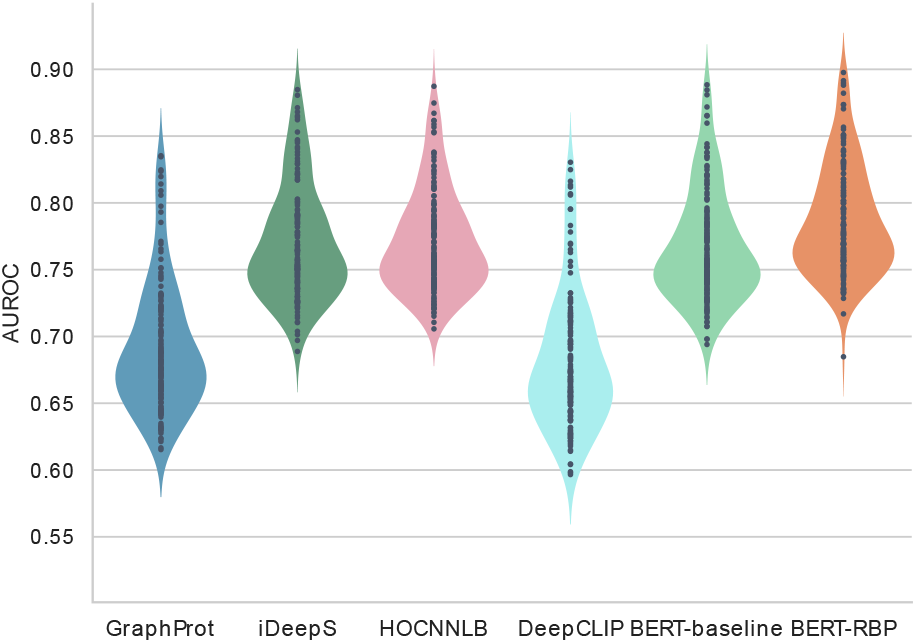
Area under the receiver operating characteristic curve (AUROC) scores of BERT-RBP and four baseline models over 154 RBP datasets. Each violin plot shows the performance of each model, and each dot within each violin plot represents the AUROC score for a single RBP dataset.

Because the original DNABERTs were pre-trained on 3-to 6-mer representations, we fine-tuned three other models, where each model takes 4-to 6-mer representations as inputs. When the AUROCs of fine-tuned models with different k-mers were compared, all fine-tuned models showed comparable performance to the 3-mer model, again demonstrating the robustness of the two-step training method (Supplementary Figure S3). The detailed comparison showed that the 3-mer model outperformed others for 135 out of 154 RBPs; thus, we refer to the 3-mer model as BERT-RBP throughout this study. In addition, since several BERT studies suggested that model performance could be improved by modifying the fine-tuning method, we evaluated two modifications of the fine-tuning architecture. One was to apply a convolution layer to all sequence embeddings after the final hidden layer, and the other was to use the weighted average of CLS tokens from all hidden layers. As a result, neither method demonstrated statistically significant improvement, suggesting that the embedding of CLS token in the final layer already had sufficient information to predict RNA-RBP interactions. (Supplementary Figure S4).

### 3.2 Attention analysis

While being a deep learning model, BERT has high interpretability (Rogers *et al.*, 2020). In this study, we investigated the types of biological information that our model could deciphered.

#### 3.2.1 Transcript Region Type

The transcript region type plays an essential role in predicting RNA-RBP interactions (Stražar *et al.*, 2016; Avsec *et al.*, 2018; Uhl *et al.*, 2020). We investigated whether the model showed high attention toward sequences from a specific transcript region, such as 5’UTR, 3’UTR, intron, or CDS, as described in section 2.4.2. The use of the CLS token stimulates the model to accumulate sequence-level information in the CLS token (Rogers *et al.*, 2020). We hypothesized that attention associated with the CLS token represents sequence-level information, that is, the transcript region type property. To test this hypothesis, we computed the relative attention to CLS associated with each region type for BERT-baseline, DNABERT, and BERT-RBP using 15 selected RBP datasets. These 15 RBPs were selected because their CLIP-seq data were analyzed and included in the previous benchmark created by Stražar *et al.* (2016). When the degree of specialization was then measured using the coefficient of variation, the relative attention level of BERT-RBP varied more than that of BERT-baseline and DNABERT for all 15 RBPs and four transcript region types (Figure 3a-d). In addition, although it had never been explicitly conditioned to collect sequence-level information to the CLS token, DNABERT still showed more variation than BERT-baseline. These results indicated that the information of the transcript region type was learned during pre-training and further enhanced during fine-tuning.

**Figure 3:**
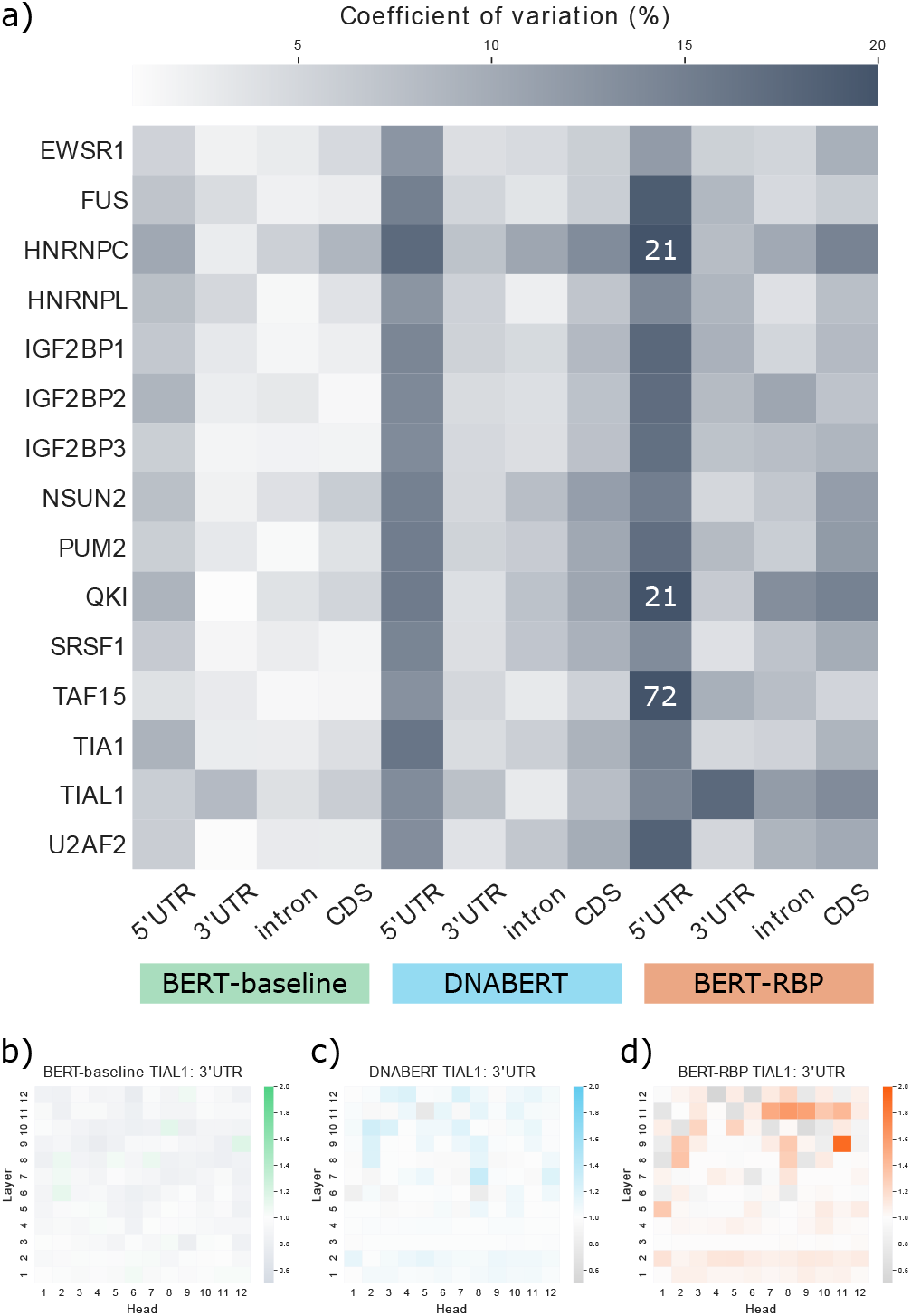
Results of sequence-level attention analysis of transcript region type. a) The degree of specialization was measured for 15 RNA-binding proteins (RBPs) and four region types using BERT-baseline, DNABERT, and BERT-RBP. The degree of specialization was evaluated using the coefficient of variation of the relative attention levels among 144 attention heads. The value was directly annotated for data points where the coefficient of variation was larger than 20%. b-d) Exemplary results of attention patterns measured by the relative attention to CLS among 144 heads. b) BERT-baseline and d) BERT-RBP trained on the same TIAL1 training set and DNABERT were analyzed using the 3’UTR annotation.

#### 3.2.2 RNA Secondary Structure

The RNA secondary structure is another feature that improves the prediction performance of several models (Maticzka *et al.*, 2014; Stražar *et al.*, 2016; Chung and Kim, 2019; Deng *et al.*, 2020). Accordingly, we investigated whether our model could consider the RNA secondary structure during prediction (Section 2.4.3). For this purpose, nine RBPs (EWSR1, FUS, hnRNPK, RBM22, SRSF1, SRSF9, TAF15, TIA1, TIAL1) with varied structural preferences were selected (Dominguez *et al.*, 2018; Adinolfi *et al.*, 2019). When DNABERT and BERT-RBP were compared, there were variations in the attention patterns for loop structures (Figure 4). Using the hnRNPK dataset, we further examined the variation of attention weights at the head 7-12, which had the highest attention ratio for the internal, hairpin, and multi-branched loops. The examination revealed a shift in specialization from DNABERT to BERT-RBP (Figure 5). The head 7-12 of DNABERT was initially specialized for detecting the dangling ends, but it began to attend more to loop structures after fine-tuning. In addition, such the shift of specialization towards loop structures was observed for the other eight RBPs (Supplementary Figure S5), and also when we analyzed BERT-RBP fine-tuned using different random seeds. These results align with the RBP’s general binding preferences toward unstructured regions (Dominguez *et al.*, 2018). Taken together, the pretrained BERT architecture can vary the type of structural information processed during fine-tuning.

**Figure 4:**
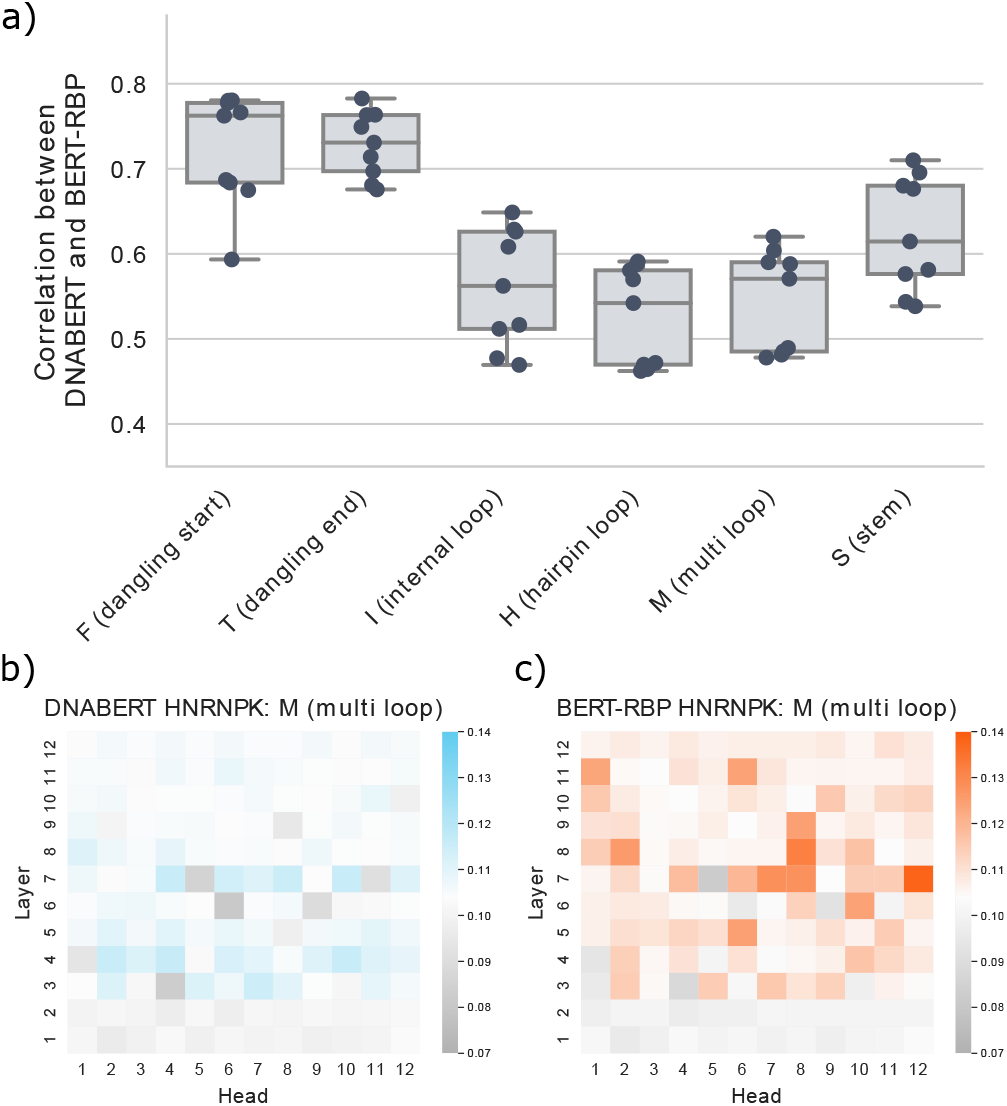
Results of token-level attention analysis of the RNA secondary structure. The similarity between DNABERT and BERT-RBP was measured for six structure types. The similarity was evaluated using Kendall’s rank correlation coefficient between the attention ratio patterns of DNABERT and BERT-RBP. For each RBP dataset, the similarity was determined for five BERT-RBPs, each of which was trained using a different random seed. Each dot represents the mean similarity score for a single RBP dataset.

**Figure 5:**
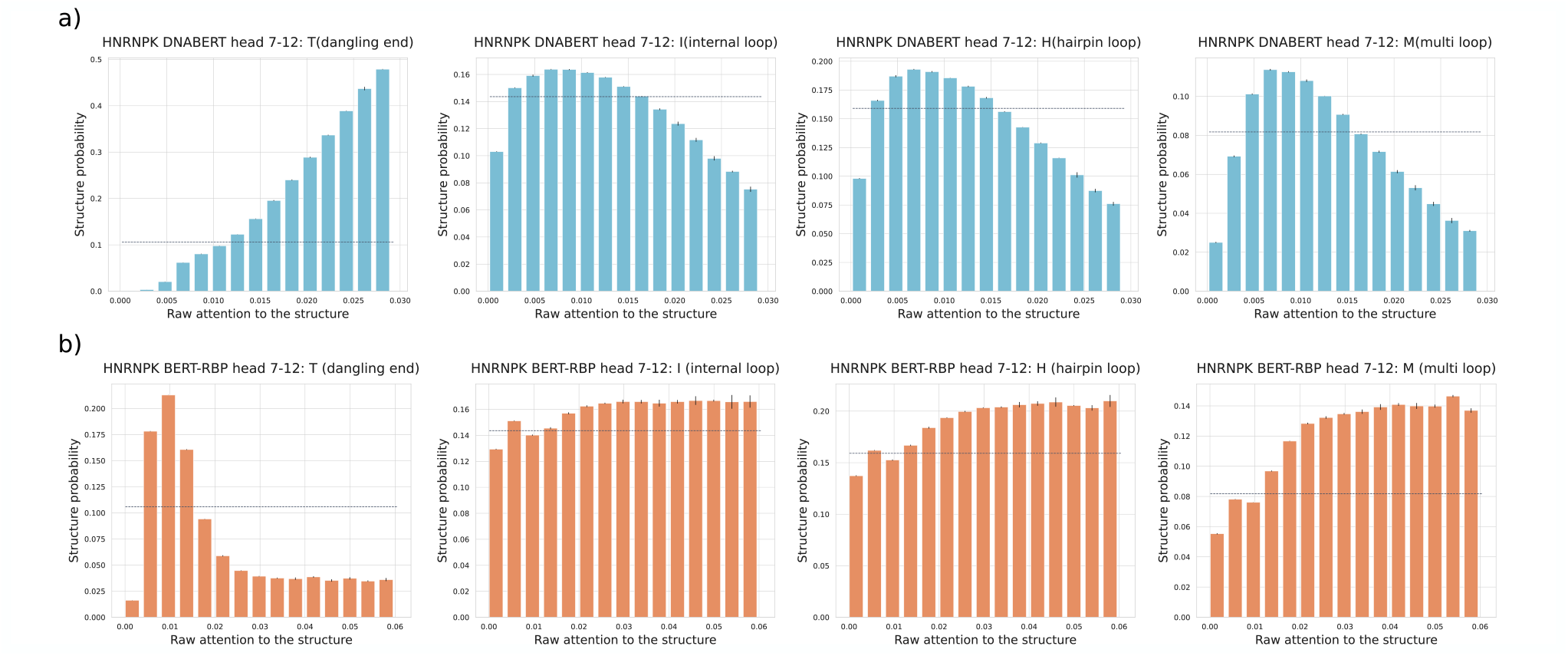
The detailed attention analysis of RNA secondary structure. a-b) Exemplary results of analysis for a) DNABERT and b) BERT-RBP using the same heterogeneous nuclear ribonucleoprotein K (hnRNPK) dataset. The relationship between the raw attention and the token probability for each structural property was measured at the head 7-12. The head was most specialized for detecting the internal, hairpin, and multi-branched loops within the BERT-RBP trained on the hnRNPK training data. The horizontal dashed lines represent the background probability of the corresponding structure within the hnRNPK non-training dataset. Error bars represent means standard deviations among three subsets randomly split from the original non-training data.

#### 3.2.3 Motif Extraction

The RBP binding motif is an essential feature of RNA sequences to distinguish sequences that bind to RBPs from others. Using high-attention regions as motif candidates, we extracted RBP binding motifs from input sequences in the test dataset (Section 2.4.4). To validate the extracted motifs, we applied an *ab initio* motif discovery algorithm, STREME, to the same dataset and compared motifs extracted from BERT-RBP to those discovered by STREME and motifs from mCrossBase. Motifs detected using BERT-RBP and STREME are listed in the Supplementary Table S3. A certain number of motifs extracted from BERT-RBP qualitatively agreed to those from STREME and mCrossBase (Supplementary Figure S6, Supplementary Table S3). There were also cases where extracted motifs did not necessarily agree, especially when the E-value calculated by STREME was relatively large. However, even using such RBP datasets, BERT-RBP still showed comparable performance to others (Supplementary Table S2); therefore, our model focused on RBP binding motifs when they were rich in information to indicate RBP binding but also used information other than sequence motifs when necessary.

## 4 Discussion

As our analysis demonstrated the model’s capability to translate transcript region type and RNA secondary structure, DNABERT can potentially be applied to other RNA-related tasks, such as RNA subcellular localization prediction (Gudenas and Wang, 2018; Yan *et al.*, 2019), RNA secondary structure prediction (Chen *et al.*, 2020; Sato *et al.*, 2021), and RNA coding potential prediction (Hill *et al.*, 2018). Another possible extension of our research is to use DNABERT to simply extract sequence features and combine extracted representations with other information such as secondary structures. It is a widely investigated topic in the field of natural language processing how to combine sequence representations from BERT with other information. Ostendorff *et al.* (2019), for instance, concatenated text representations from pre-trained BERT with separately embedded metadata and author information to classify books. When using a BERT model as the feature extractor, it would be reasonable to compare the quality with other methods to embed sequences into distributed representations, such as word2vec (Mikolov *et al.*, 2013a,b) and embeddings from language models (ELMo) (Peters *et al.*, 2018).

Although our analysis implied that the fine-tuned model could utilize the information learned by DNABERT, it must be noted that the attention analysis itself does not necessarily provide a comprehensive explanation of how the model processes information. (Vig *et al.*, 2021; Jain and Wallace, 2019) Diagnostic probing is a method to train a simple linear regression model by using attention weights or hidden states as inputs and labeled information as targets (Liu *et al.*, 2019). However, it should be noted that probing is an indirect method to analyze information with additional training using highdimensional vectors. Abnar and Zuidema (2020) proposed attention flow to quantify the cumulative amount of attention from the final embeddings to the first input tokens, but this method requires *O*(*d*^2^ * *n*^4^) computation, where *d* is the number of Transformer layers, and *n* is the number of tokens. The syntactic relationship among tokens is another intensely researched topic in BERTology and may incorporate hidden contextual patterns of nucleotide sequences (Goldberg, 2019). In the context of protein BERT models, it was recently demonstrated that protein contact maps could be reconstructed using the attention maps extracted from the pre-trained protein BERT model (Rao *et al.*, 2021). If one could overcome the difference in the frequency of tokens in contact, it would be possible to reconstruct the base-pairing probability matrix using the attention maps of a nucleotide BERT model. There is yet no single attention analysis method to provide a comprehensive explanation, but combining the above methods may complement each other.

In this study, we proposed BERT-RBP, a fine-tuned BERT model for predicting RNA-RBP interactions. Using the eCLIP-seq data of 154 different RBPs, our model out-performed state-of-the-art methods and the baseline BERT model. Attention analysis revealed that BERT-RBP could distinguish both the transcript region type and RNA secondary structure using only sequence information as inputs. The results also inferred that the attention heads of BERT-RBP could either utilize information acquired during DNABERT pre-training and vary the type of information processed when necessary. Thus, this study provides a state-of-the-art tool to predict RNA-RBP interactions and infers that the same method can be applied to other bioinformatics tasks.

## Acknowledgements

Computations were partially performed on the NIG supercomputer at ROIS National Institute of Genetics. We acknowledge Dr. Shitao Zhao and Taro Matsutani for their support and comments on the manuscript.

## Funding

This work was supported by the Ministry of Education, Culture, Sports, Science, and Technology (KAKENHI) [grant numbers: JP17K20032, JP16H05879, JP16H06279, and JP20H00624 to MH] and JST CREST [grant numbers: JPMJCR1881 and JPMJCR21F1 to MH].

## Conflict of Interest

none declared.

## Supplementary data

Supplementary data including Figures S1, S2, S3, S4, S5, and S6, and Tables S1, S2, and S3 are available.

## Notes

### Competing Interest Statement

The authors have declared no competing interest.

### Summary of Updates

(Mar 2022): Paper throughly revised. Main changes: (1) updated the results comparing model performance (2) updated attention analysis results regarding probabilistic behavior of model training (3) added motif analysis

