## Supplementary Material for "Prediction of RNA-protein interactions using a nucleotide language model"

### Supplementary materials for "Prediction of RNA-protein interactions using a nucleotide language model"

Supplementary Table S 1: Hyperparamaters used during the training of BERT models

| Hyperparameter | BERT-baseline | BERT-RBP | BERT-RBP(with CNN) | BERT-RBP(CLS average) |
|---|---|---|---|---|
| Batch size | 64 | 64 | 64 | 64 |
| Learning rate | 6e-5 | 2e-4 | 2e-4 | 2e-4 |
| Epoch | 5 | 3 | 3 | 3 |
| Warmup rate | 0.1 | 0.1 | 0.1 | 0.1 |
| Dropout probability | 0.01 | 0.1 | 0.01 | 0.01 |
| Weight decay rate | 0.01 | 0.01 | 0.01 | 0.01 |

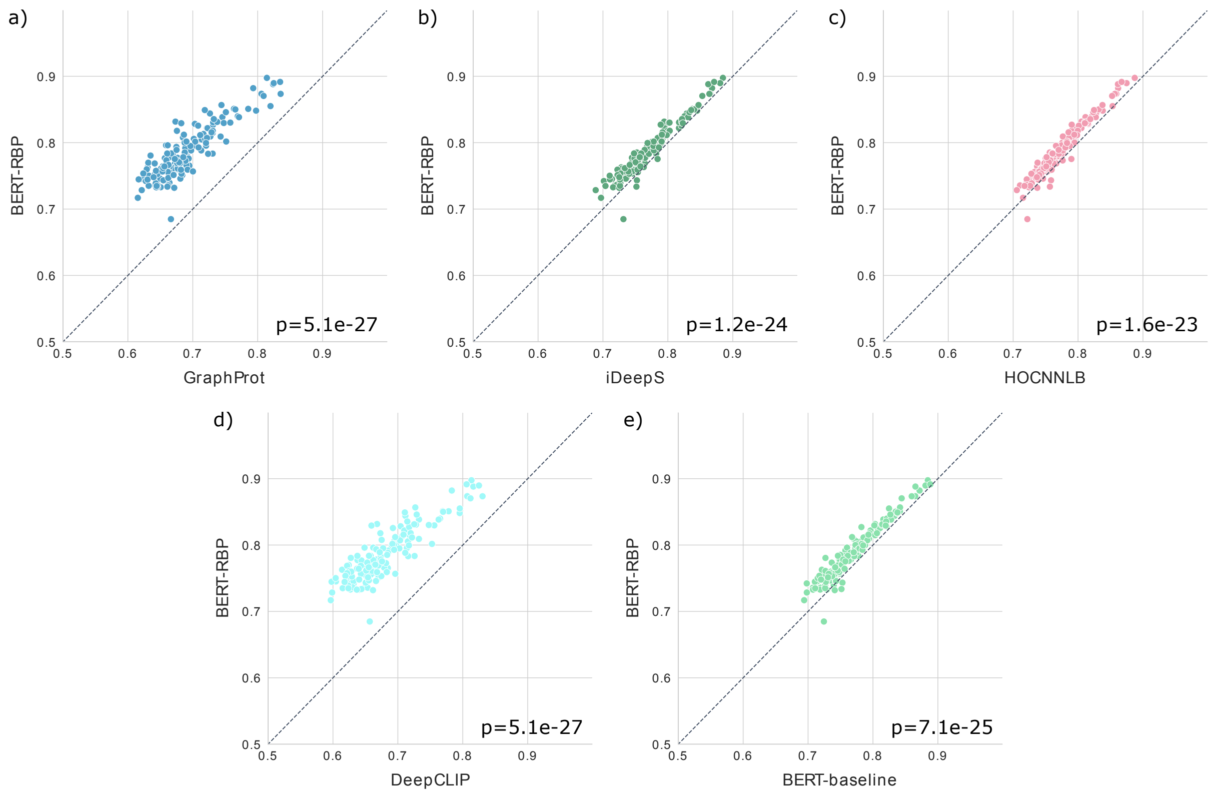

Supplementary Figure S 1: Detailed comparison of BERT-RBP's performance against a) GraphProt (Maticzka *et al.*, 2014), b) iDeepS (Pan *et al.*, 2018), c) HOCNNLB (Zhang *et al.*, 2019), d) DeepCLIP (Grønning *et al.*, 2020), or e) BERT-baseline by the measure of AUROC. Each dot represents the AUROC scores of BERT-RBP and the corresponding baseline model trained using the same RBP dataset. The diagonal dashed line indicates where the performances of the two models are identical. p-values were calculated using Wilcoxon's signed-rank test.

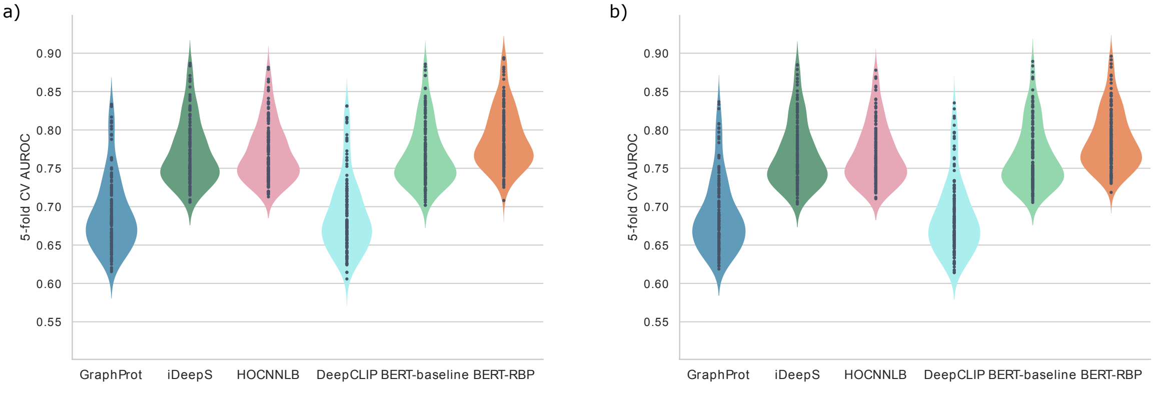

Supplementary Figure S 2: Cross-validation performance was measured using area under the receiver operating characteristic curve (AUROC) scores for BERT-RBP and four baseline models over 154 RBP datasets. Each violin plot shows the performance of each model, and each dot within each violin plot represents the AUROC score for a single RBP dataset. Training and evaluation datasets were used for a), and low sequence-similarity datasets were used for b).

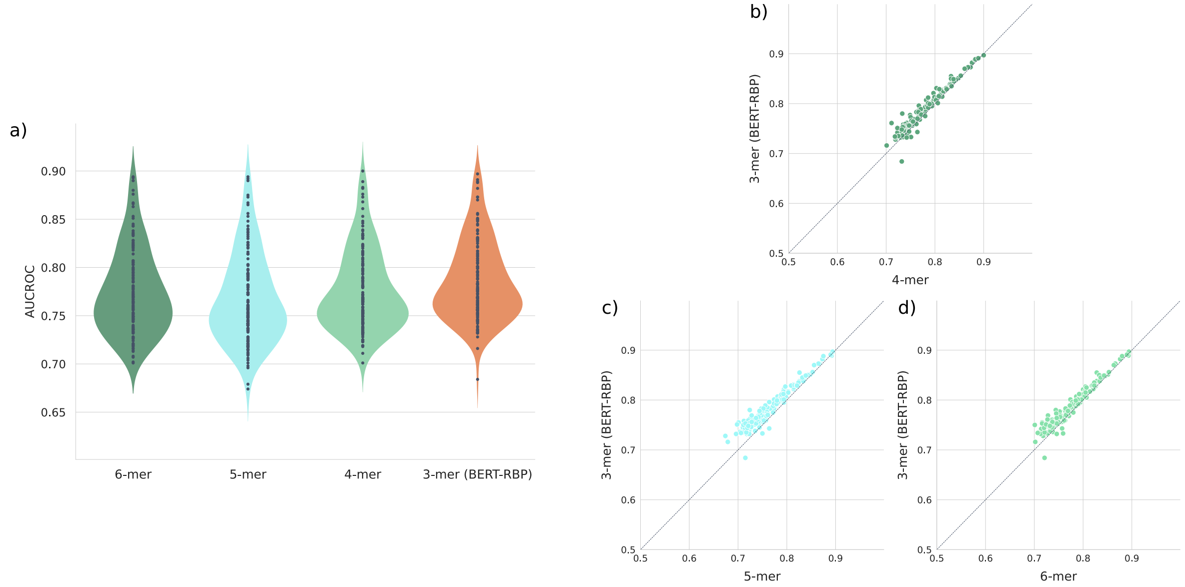

Supplementary Figure S 3: Comparison of different k-mer models. a) Area under the receiver operating characteristic curve (AUROC) scores of models pre-trained and fine-tuned with different k-mer representations in over 154 RNA-binding protein (RBP) datasets. Each violin plot shows the performance of each model, and each dot within the violin plot represents the AUROC score for a single RBP dataset. b-d) A detailed comparison of BERT-RBP's performance against b) 6-mer, c) 5-mer, or d) 4-mer models by AUROC measurement. Each dot represents the AUROC scores of BERT-RBP and the target model trained using the same RBP dataset. The diagonal dashed line indicates that the performances of the two models are identical.

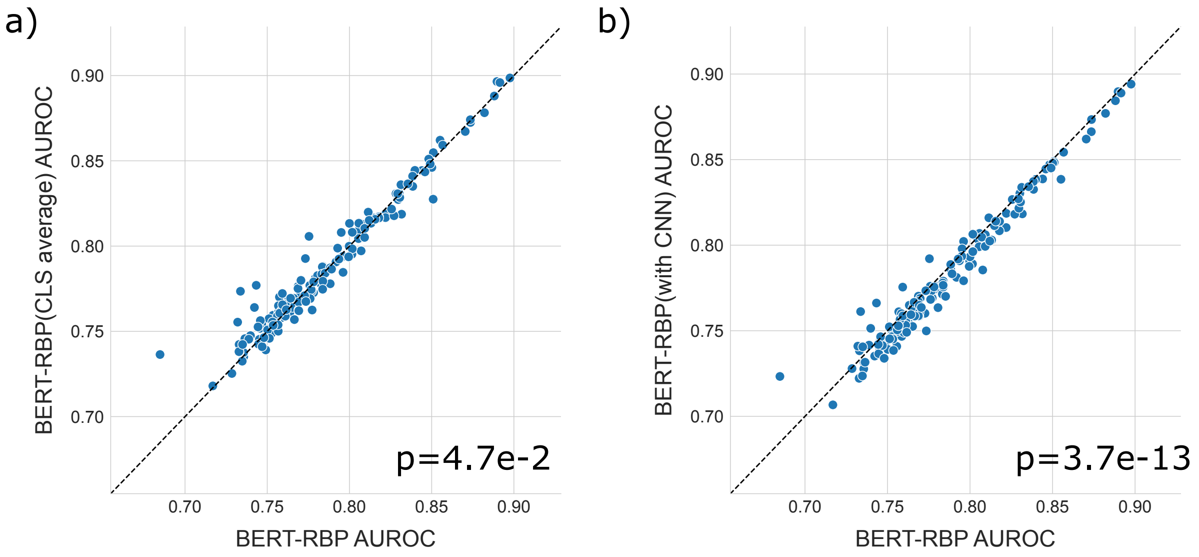

Supplementary Figure S 4: Detailed comparison of BERT-RBP's performance against a) BERT-RBP with a convolution layer after the final layer and b) BERT-RBP that uses the weighted average of CLS tokens from all hidden layers. Each dot represents the AUROC scores of BERT-RBP and the target model trained using the same RBP dataset. The diagonal dashed line indicates that the performances of the two models are identical. p-values were calculated using Wilcoxon's signed-rank test.

### EWSR1

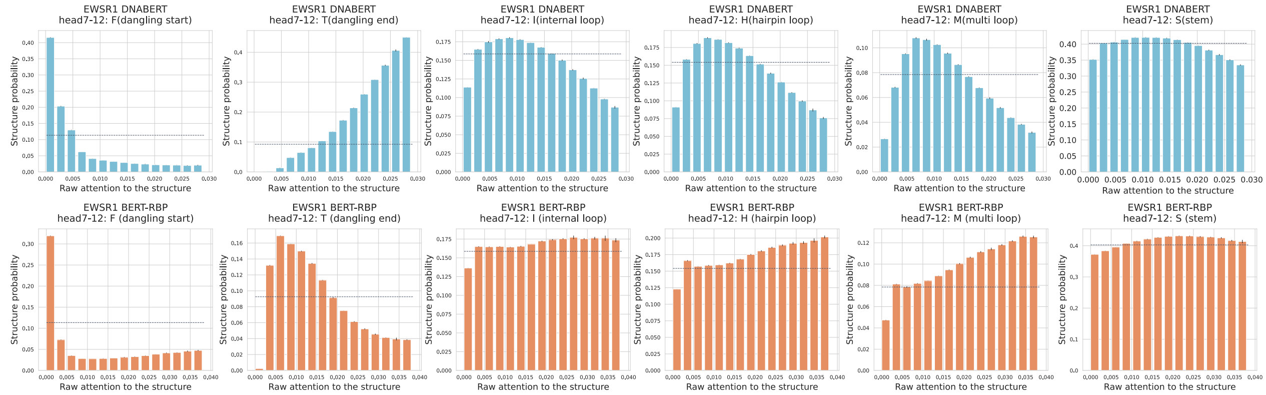

### FUS

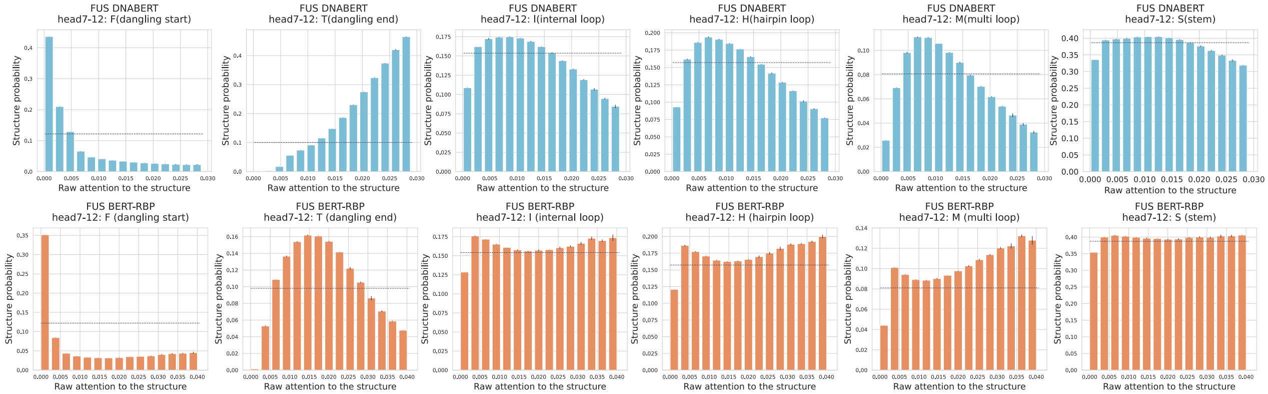

### HNRNPK

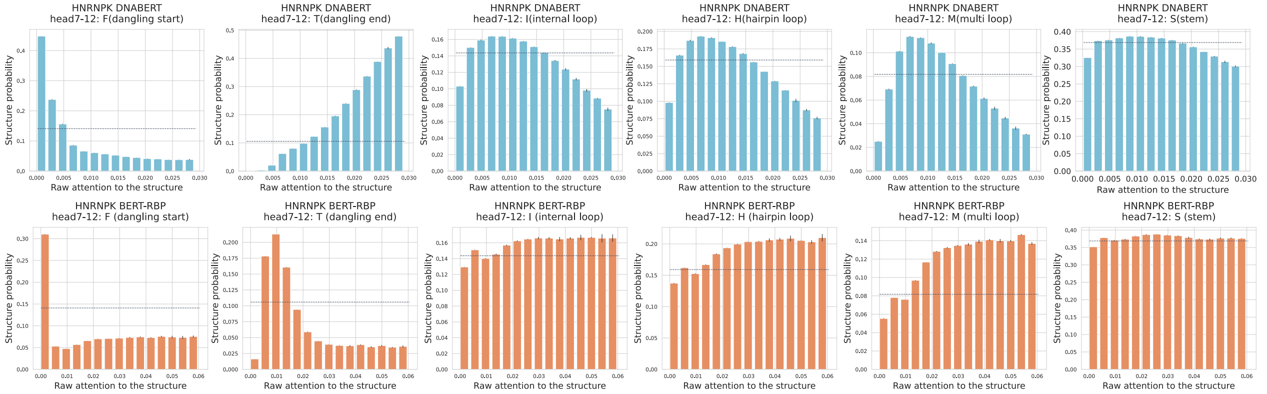

### RBM22

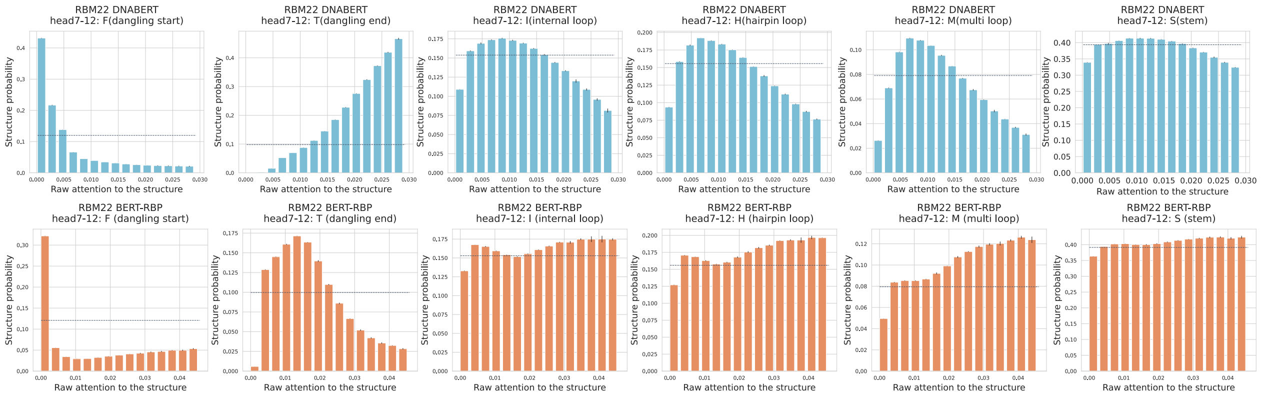

### SRSF1

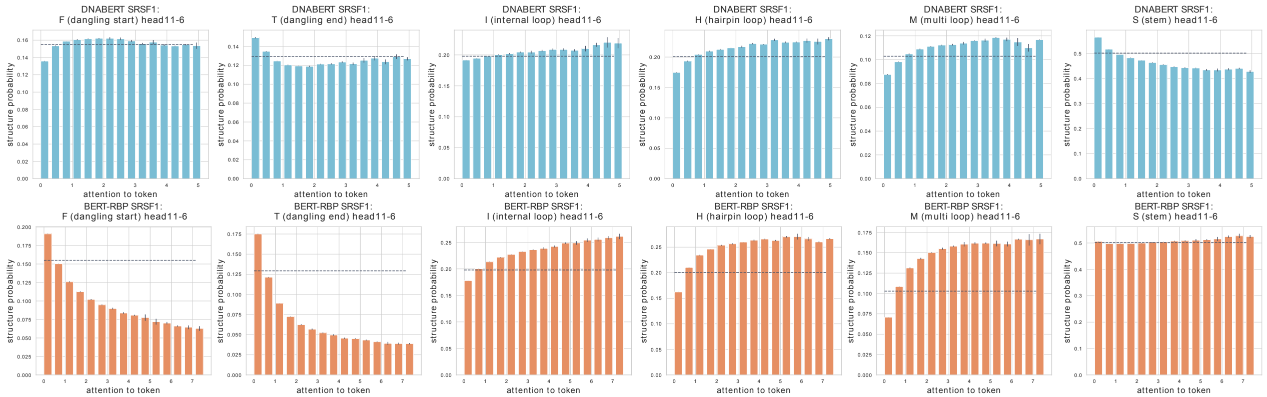

### SRSF9

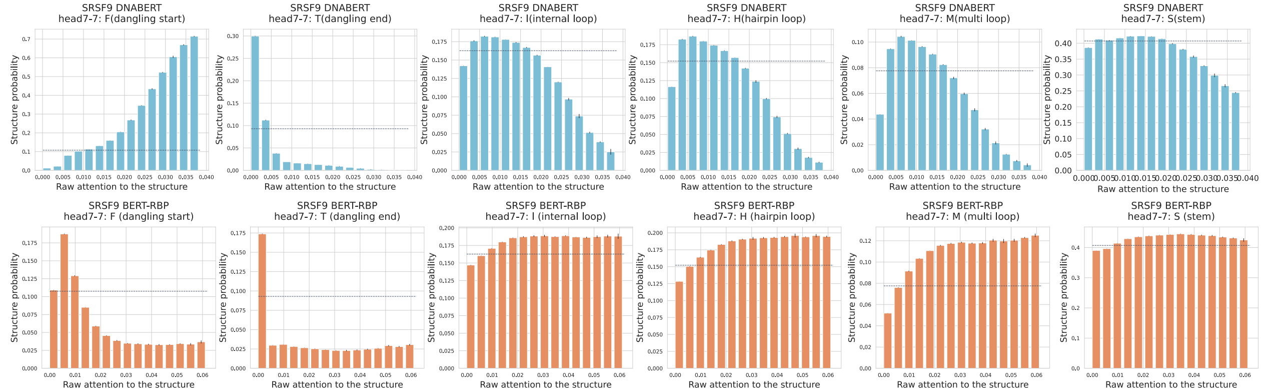

### TAF15

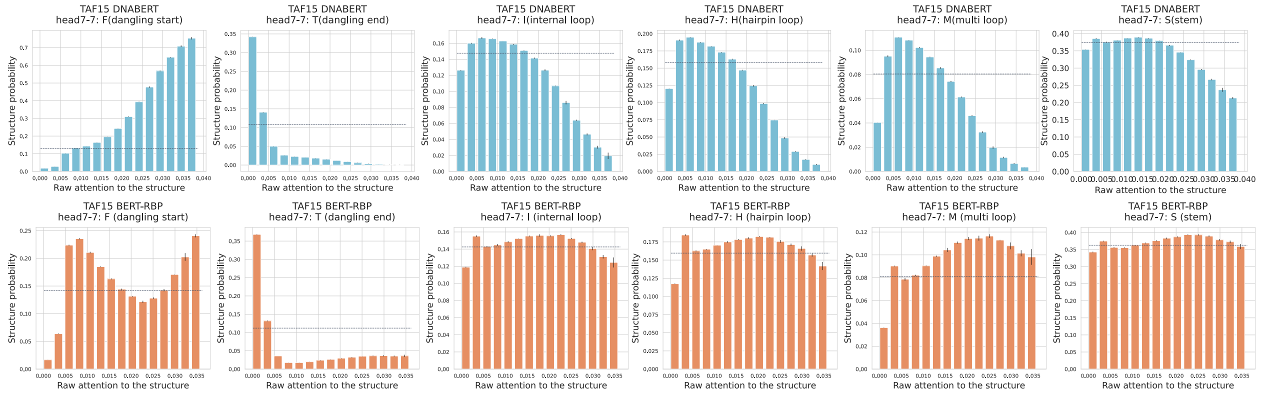

### TIA1

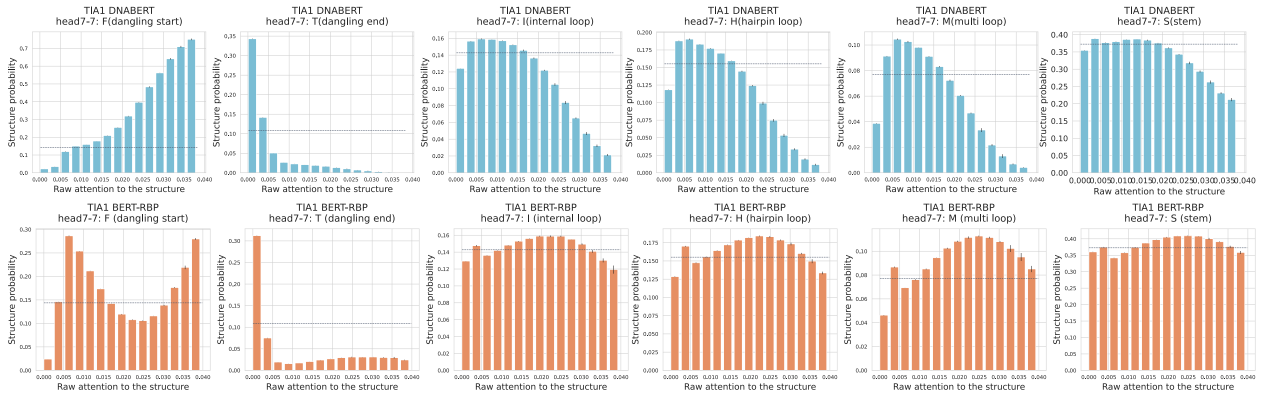

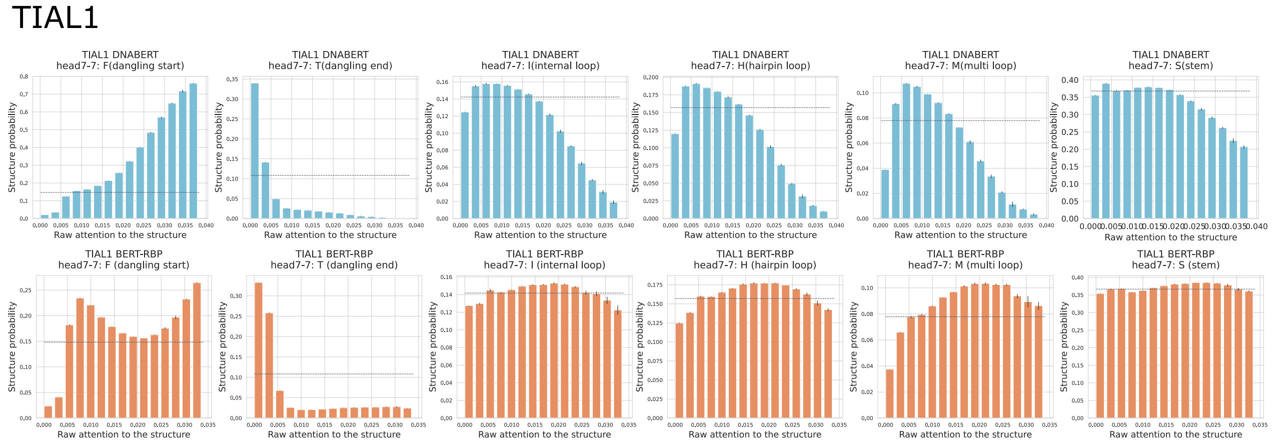

Supplementary Figure S 5: The detailed results of correlation analysis for nine RNA-binding proteins (RBPs) that showed the shift of specialization. A detailed analysis was conducted to inspect correlations between the raw attention to the RNA secondary structure type and the token probability to incorporate the structure at the selected head. For each BERT-RBP, the attention head with the highest attention ratio to the structure was chosen (bottom, orange), and the head of DNABERT (Ji *et al.*, 2021) at the same position was used (top, blue). The horizontal dashed lines represent the background probability of the structure label within each RBP non-training dataset. Error bars represent means ± standard deviations among three subsets randomly split from the original non-training set.

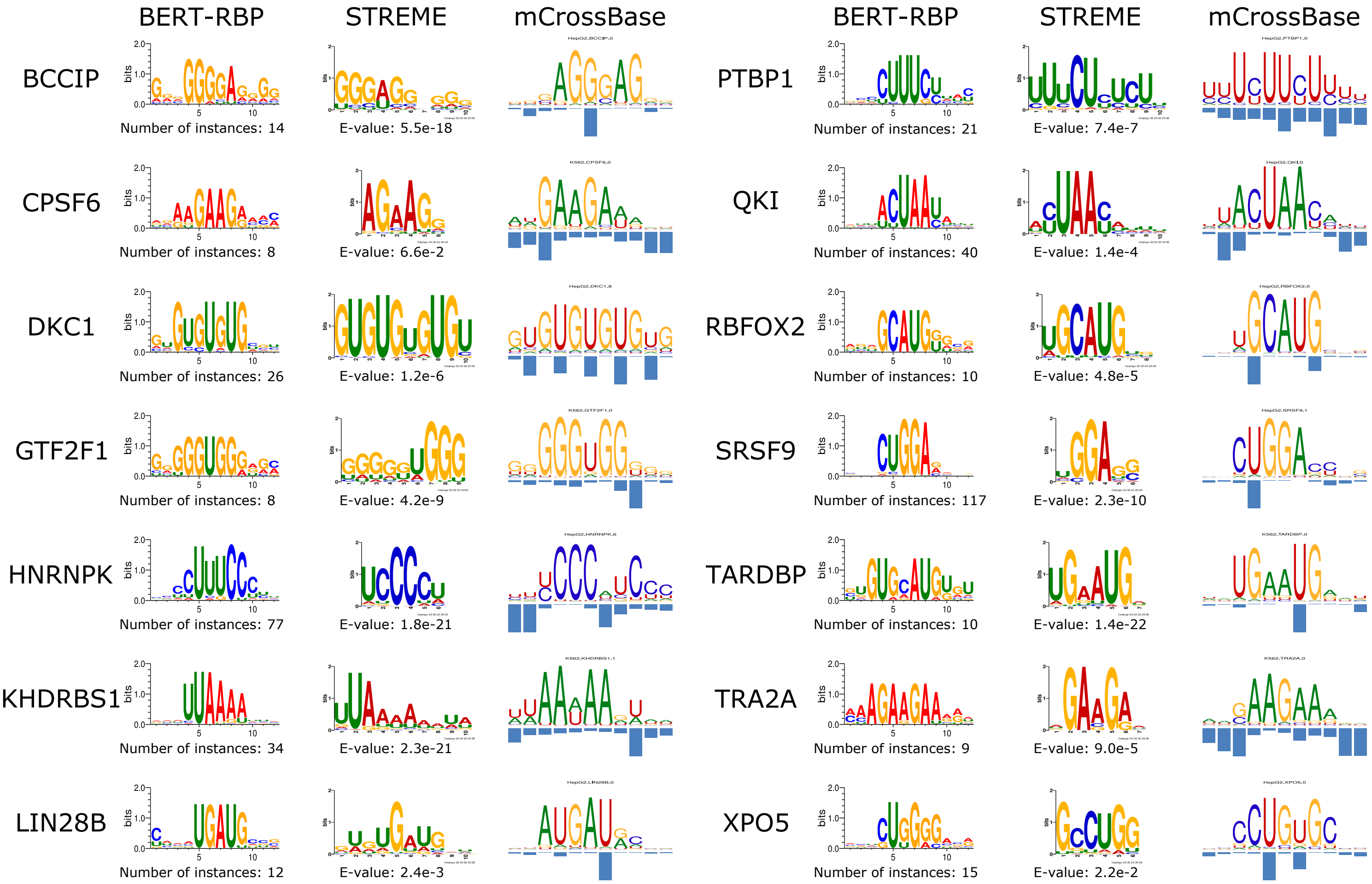

Supplementary Figure S 6: Exemplary motifs extracted from BERT-RBP that agree to motifs detected by STREME (Bailey, 2021) and motifs downloaded from mCrossBase (Feng *et al.*, 2019).
